# Microbial Metagenomes Across a Complete Phytoplankton Bloom Cycle: High-Resolution Sampling Every 4 Hours Over 22 Days

**DOI:** 10.1101/2024.09.27.614549

**Authors:** Brook L. Nunn, Emma Timmins-Schiffman, Miranda C. Mudge, Deanna Plubell, Gabriella Chebli, Julia Kubanek, Michael Riffle, William S. Noble, Elizabeth Harvey, Tasman A. Nunn, Marcel Huntemann, Alicia Clum, Brian Foster, Bryce Foster, Simon Roux, Krishnaveni Palaniappan, Supratim Mukherjee, T.B.K. Reddy, Chris Daum, Alex Copeland, I-Min A. Chen, Natalia N. Ivanova, Nikos C. Kyrpides, Tijana Glavina del Rio, Emiley A. Eloe-Fadrosh

## Abstract

In May and June of 2021, marine microbial samples were collected for DNA sequencing in East Sound, WA, USA every 4 hours for 22 days. This high temporal resolution sampling effort captured the last 3 days of a *Rhizosolenia* sp. bloom, the initiation and complete bloom cycle of Chaetoceros socialis (8 days), and the following bacterial bloom (2 days). Metagenomes were completed on the time series, and the dataset includes 128 size-fractionated microbial samples (0.22-1.2 *µ*m), providing gene abundances for the dominant members of bacteria, archaea, and viruses. This dataset also has time-matched nutrient analyses, flow cytometry data, and physical parameters of the environment at a single point of sampling within a coastal ecosystem that experiences regular bloom events, facilitating a range of modeling efforts that can be leveraged to understand microbial community structure and their influences on the growth, maintenance, and senescence of phytoplankton blooms.

## Background and Summary

Phytoplankton blooms, from initiation to decline, are crucial for biogeochemical cycling. They absorb carbon dioxide and nutrients from surface waters^1^, fueling the trophic structure in the euphotic zone and influencing secondary production^2^. Additionally, as these blooms sink, they contribute to carbon sequestration at depth^3^. Nutrient and light conditions are traditionally thought to drive the timing and intensity of phytoplankton blooms^4^ but decades of research have revealed that physical and chemical interactions between prokaryotic community members can significantly modulate eukaryotic blooms^5,6^. These cross-kingdom interactions are dynamic and can be influenced by environmental variables, controlling the rise and fall of different species and driving biogeochemical cycles in coastal environments and in the open ocean. The transfer of biotic and abiotic chemicals among many interacting organisms can influence phytoplankton’s growth, maintenance, and senescence, influencing large-scale events like blooms^5^. These complex community relationships have been monitored at different time intervals, from weekly^7,8^ to daily^9^ across phytoplankton bloom events. For example, a study across 3 days on a 4-hour time scale revealed how heterotrophic bacterial gene expression is diel in nature, trailing in timing behind phytoplankton gene expression^10^. This emphasized unforeseen, rapid responses between oceanic prokaryotes and eukaryotes that have not yet been captured over the full course of bloom events. The current dataset fills a critical gap by providing high temporal resolution field collections across multiple bloom cycles, providing ecosystem-level details that hold the potential to inform community-level controls on the initiation, maintenance, and termination of phytoplankton blooms.

In May and June of 2021, we sampled coastal waters in East Sound, WA, by isolating the microbial size classes (0.22-1.2 *µ*m) for metagenomic analyses *every 4 hours for 22 days*. East Sound, WA is a fjord enveloped by Orcas Island with 4500 coastal residents and is important to local tourism, fishing, and shellfishing. This dynamic coastal ecosystem experiences a tidal variation of up to 11.6 ft (Figure 1A)^11^ and is influenced by terrestrial runoff. East Sound was chosen due to its economic and social significance and because it is impacted annually by algal blooms. A fluorescent probe mounted at the intake for all samples at 2 m deep in the water revealed that chlorophyll *a* concentrations peaked on June 4, 2021, at 94 *µ*g L^-1^ due to an intense *Chaetoceros socialis* bloom that lasted 8 days (Figure 1B; Figure S1)^12^. This bloom was followed by nine days of low chlorophyll fluorescence. In the final days of the sampling effort, the 3-day initiation period of a second *C. socialis* bloom was captured.

**Figure 1.**
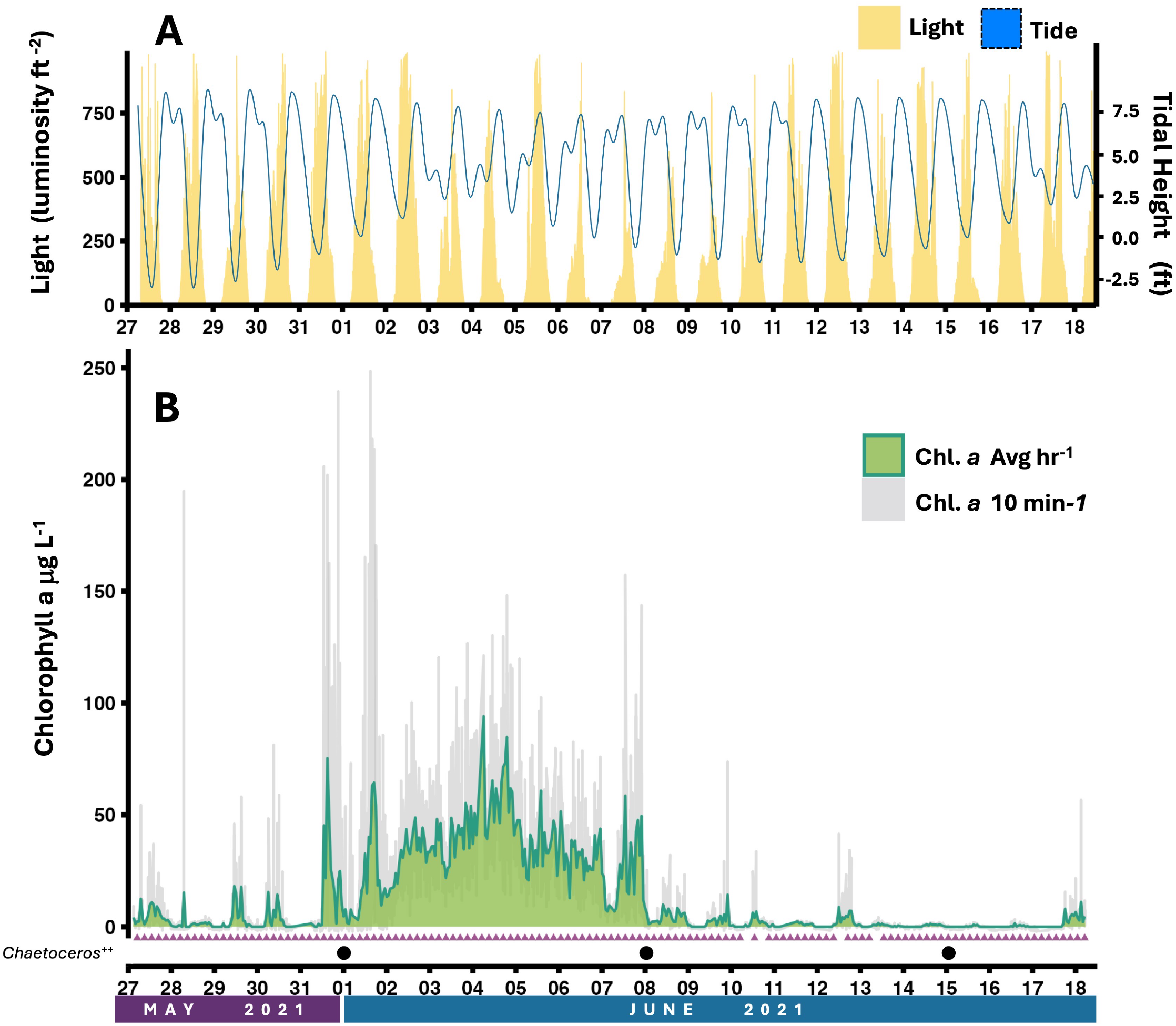
**A.** Light levels at 2 m depth at the sampling site measured with HOBO data logger (yellow) and the variation in tidal height from NOAA station 9449771 over the 22 days (blue)^11^. **B.** Direct fluorometry measurements were collected at 10-minute intervals using a YSI probe (grey line) and converted to chlorophyll a (chl a) concentrations. The green line represents chl a concentrations estimated through binning data by hour^20^. Purple triangles denote time points where DNA was sequenced successfully. DNA was collected 6 times per day at 1:00, 5:00, 9:00, 13:00, 17:00, and 21:00. Missing triangles in the time series did not have sufficient DNA for sequencing (5 time points). Black dots denote time points where SoundToxins confirmed *Chaetoceros spp*. was the dominant phytoplankton in water. SoundToxins also confirmed *Chaetoceros spp*. dominance on May 25, 2021 and June 22, 2021.

Samples were collected near Orcas Island in East Sound, WA from May 27 (13:00) to June 18 of 2021 (13:00) 100 m offshore and 2 m below the surface. Time-matched microbial metagenomes (0.22-1.2 *µ*m), nutrient analyses, flow cytometry data, and coastal physical parameters are provided here.

## Methods

### Sampling protocol

Samples were collected near Orcas Island in East Sound, WA from May 27 (13:00) to June 18 (13:00) of 2021. Cross-linked polyethylene tubing (PEX) with 0.5 inch diameter was fixed to a 3-point buoy moored 100 m offshore (Lat: N 48° 40.590’, Long: W 122° 52.994’) with a set intake at 2 m below the surface, to allow for tidal fluctuations. Water was directly pumped to a temporary field station using a Water Diaphragm Pump (SEAFLO) with a rheostat set to low to decrease the flow rate to 5 L min^-1^. Water collection volumes were monitored at each timepoint to as an observational calibration to ensure consistency of the SEAFLO pump across the field season. Incoming waters were pulled through a 15 cm diameter funnel with 200 *µ*m nylon mesh. Samples were collected every 4 hours. Prior to collecting each time point, the 100 m of PEX transfer tubing was rinsed with three tubing volumes of the seawater at that time. Each sample type had a designated collection carboy; the carboys were triple-rinsed with incoming seawater before each collected time point. Whole water is defined as water filtered only through the 200 *µ*m mesh at the collection funnel and was used for nutrient analyses and flow cytometry. A valve was used to alternatively pass the incoming 200 *µ*m filtered water through a series of 3 string filters 100 *µ*m, 10 *µ*m, 1 *µ*m (Hydronix SWC Universal Whole House String Wound Sediment Water Filter Cartridge 2.5” x 10”) yielding pre-filtered water for downstream microbiome collections. Below are the details of the specific sample collection types per time point. Samples were always collected in the following order (parenthetical numbers indicate biological replicates): nutrients, flow cytometry, prokaryotic DNA (1), prokaryotic proteomic (5), eukaryotic DNA/RNA (3), eukaryotic proteomics (1), and dissolved metabolites (3). Here, we provide all datasets that have been analyzed and quality-controlled thus far; this includes the environmental metadata, nutrients, flow cytometry, and prokaryotic DNA sequences.

### Environmental metadata

A HOBO data logger (HOBO Pendent MX Temperature/Light Data Logger) was also deployed at the site to conduct constant measures of depth, temperature, and light (luminosity/ft^2^) (Figure 1A; Dataset 1). Weather and tidal data for the dates of May 27-June 18, 2021 were downloaded from the National Oceanic and Atmospheric Administration (NOAA) Tides and Currents portal for Rosario, Orcas Island, WA, Station ID: 9449771 as reported using a harmonic prediction type^11^.

An EXO1 Multiparameter Sonde (YSI) was deployed at the collection site at a constant depth of 2 m and set to collect *in situ* measurements every 10 minutes. The probe had four ports which were fitted with sensors (YSI) to measure conductivity/temperature (YSI EXO Conductivity and Temperature Smart Sensor, SKU 599870), dissolved oxygen (EXO Optical Dissolved Oxygen Smart Sensor, SKU 599100-01), pH (EXO pH Smart Sensor, SKU 577601), and Total Algae (Chlorophyll, Phycocyanin, Phycoerythrin; EXO Total Algae PE Smart Sensor, SKU 599103-01)^12^. Data from the probe was manually downloaded each day at 14:00 before redeployment and the probe was calibrated every 5 days. The Total Algae sensor covers a range of 0.1 to 400 *µ*g L^-1^ (0 to 100 RFU), with a detection limit of ∼0.1 *µ*g L^-1^, and a resolution of 0.1 *µ*g L^-1^ Chl (0.1% RFU). Total algae measurements taken every 10 minutes were plotted as chlorophyll *a* concentrations in *µ*g L^-1^ (binned and averaged by the hour; Figure 1B)^12^. Weekly relative abundance measures of phytoplankton at the East Sound monitoring site by the SoundToxins Phytoplankton Monitoring Network, Washington Sea Grant program, confirm the presence of Rhizosolenia at the beginning of the field collection and *Chaetoceros spp*. throughout the sample collection (Figure 1B; Dataset 2)^12^.

### Nutrients

Water samples for nutrients were collected by rinsing a 1 L Pyrex media bottle and a 60 mL HDPE syringe in triplicate with whole water. From a fresh collection of 1 L, we filtered 50 mL whole water using the cleaned syringe through a 25 mm 0.45 *µ*m cellulose syringe filter (Nalgene surfactant-free). Filtered water samples were collected in triple-rinsed 50mL HDPE bottles and stored immediately at -20 °C. Bottles were transferred to the University of Washington for long-term storage at -20 °C. Total nutrient analysis was performed in triplicate by the University of Washington Marine Chemistry Lab to determine concentrations of nitrate, nitrite, ammonium, silicate, and phosphate following standard methods outlined in UNESCO, 1994^13,14^ (Figure 2; Dataset 3)^12^.

**Figure 2.**
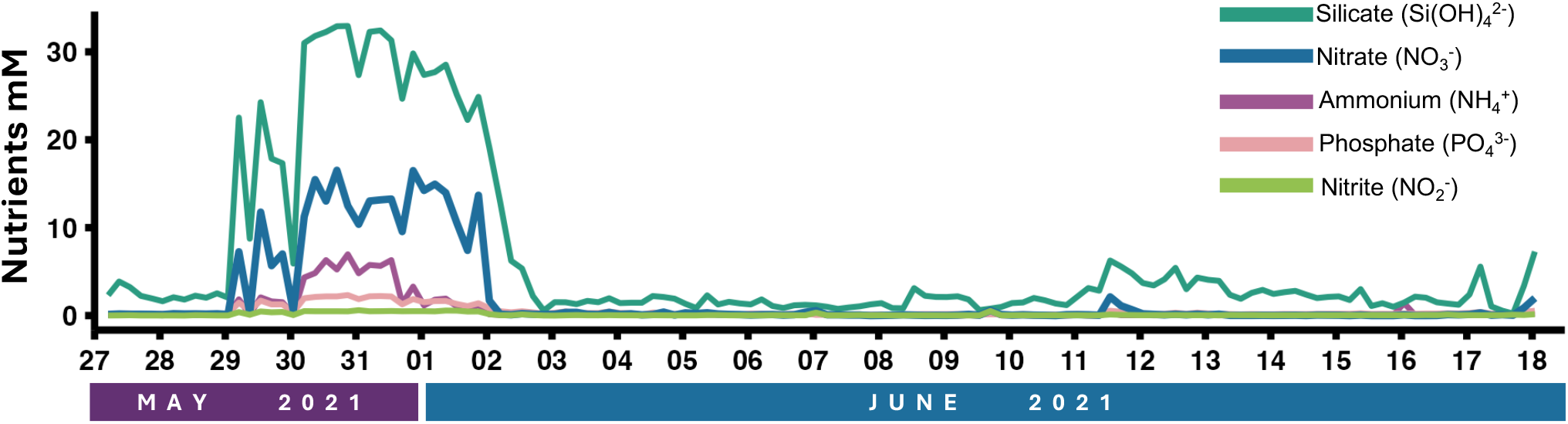
Nutrient concentrations (mM) of silicate (teal), nitrate (blue), ammonium (purple), phosphate (pink), and nitrite (lime green) across the 22 day timeframe. Samples were collected at 2 m depth at 1:00, 5:00, 9:00, 13:00, 17:00, and 21:00 each day at the East Sound sampling site.

### Flow cytometry

Two samples (one bacterial, one phytoplankton) were collected for downstream flow cytometry analyses at each time point. The bacterial fraction was specifically isolated for flow cytometry by collecting 1 mL of whole water from a glass bottle rinsed three times with incoming seawater. The 1 mL aliquot was transferred to a cryovial and immediately fixed with 20 *µ*L glutaraldehyde (25% EM-grade, MilliporeSigma) at 4 °C for 25 minutes and flash-frozen in liquid nitrogen in the field prior to long-term storage in the lab at -80 °C. For flow cytometry analysis of the phytoplankton fraction, whole water (3.5 mL) was subsampled from the same triple rinsed glass bottle and then fixed with 100 uL formaline: hexamine (18%:10% v/w) at 4°C for 25 minutes. The sample was immediately frozen in liquid nitrogen in the field prior to long-term storage in the lab at -80°C.

Each sample was analyzed on a Guava EasyCyte HT (Luminex) flow cytometer using the native software package Incyte. For bacteria, 190 *µ*L of each sample was stained with 10 *µ*L of 200X SYBR Green I (Invitrogen, Catalog number S7563) for 30 min in the dark at room temperature and distinguished based on plots of forward scatter and green fluorescence (512 nm). Biological duplicates were analyzed and averages and standard deviations were calculated (Figure 3A; Dataset 4). For phytoplankton, three major phytoplankton groups (cyanobacteria, picoeukaryotes (2-5 microns), and nanoeukaryotes (5-10 microns)) were distinguished by size based on plots of forward scatter vs. red fluorescence (692 nm). Predefined gates for each group were set up based on forward scatter and red fluorescense determined using variable size beads to calculate the forward scatter. The cyanobacteria is further distinguished by gates that target cells that have lower forward scatter with lower chlorophyll and higher orange fluorescence (phycoerythrin) relative to the picoeukaryotes. Biological triplicates were analyzed, measurements were averaged and standard deviations were calculated (Figure 3 B-D; Dataset 4)^12^.

**Figure 3.**
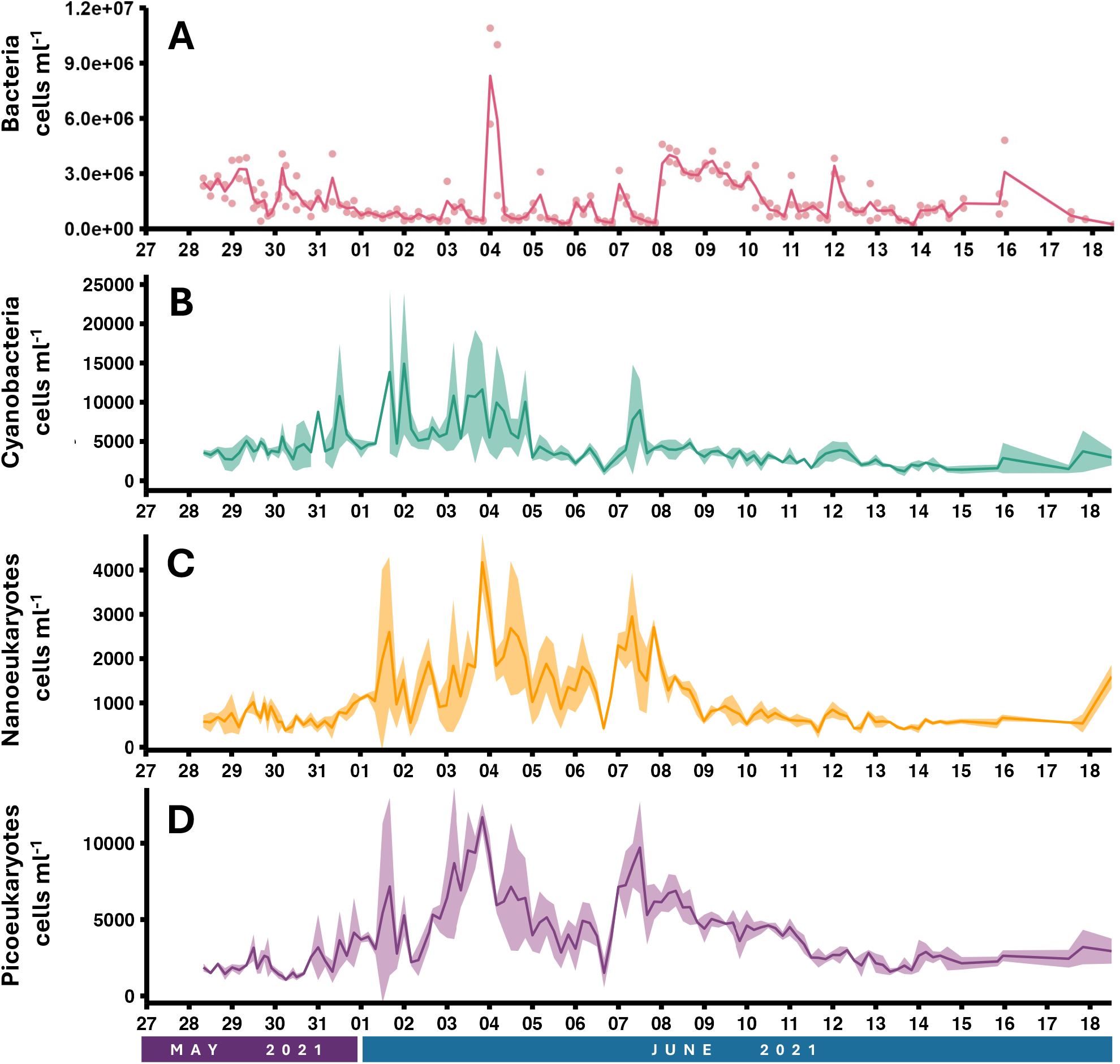
Average cell counts for **A.** bacteria in duplicates (pink) and **B.** cyanobacteria (teal), **C.** nanoeukaryotes (yellow), and **D.** picoeukaryotes (purple) in triplicate plotted over the 22 day time course. Samples were collected at 1:00, 5:00, 9:00, 13:00, 17:00, and 21:00 each day. Cell count samples were collected at a depth of 2 m at the East Sound field site. Line indicates the average for either duplicate or triplicate measurements. Individual points included for duplicate measurements and shading represents the standard deviation for triplicate measurements.

### Microbiome sample collection for metagenomics

The 1-2 L of pre-filtered water was collected and then passed through a 47 mm × 0.22 *µ*m polyethersulfone filter (Tisch, part number SF15021) using the Alexis peristaltic field pump (Proactive Environmental) at a flow rate of <0.1 L min^-1^ to isolate the bacterial fraction. Filters were immediately placed in plastic resealable baggies (5 × 4 cm), then 200 *µ*l of DNA/RNA Shield (Zymo Research) was added to the whole filter within 1 minute of collection and filter baggies were transferred to liquid nitrogen while in the field. Samples were transferred to the University of Washington and placed in a -80°C freezer prior to extraction.

### Microbiome DNA Extraction

DNA extraction was performed using Qiagen’s DNeasy PowerWater kit following the manufacturer’s instructions. Quant-iT DNA Assay Kits (Invitrogen) were used to generate accurate DNA quantitation (at the recommended molecular weight) in conjunction with our fluorescent plate reader (Varioskan Lux, SkanIt Software 7.0.2). On average, 1.3 *µ*g of high-quality DNA was isolated from each time point collected and 300 ng of isolated DNA was aliquoted into DNA-free tubes (Matrix, catalog #3743) and sent to the Department of Energy (DOE) Joint Genome Institute (JGI) for sequencing.

### Metagenome sequencing and analysis

Of the 133 time points collected between May 27 (13:00) to June 18 (13:00) of 2021, a total of 128 DNA samples were successfully sequenced at the Department of Energy (DOE) Joint Genome Institute (JGI) using Illumina technology. The five samples that did not have adequate DNA included 6/10/2021 17:00, 6/11/2021 01:00, 6/12/2021 21:00, 6/13/2021 01:00, and 6/18/2021 13:00.

#### Library Creation

Plate-based DNA library preparation for Illumina sequencing was performed on the Hamilton Vantage robotic liquid handling system using the standard Hamilton Run Control software and with the Kapa Biosystems HyperPrep library preparation kit (Roche). 200 ng of genomic sample DNA was sheared to 600 bp using a Covaris LE220 focused-ultrasonicator for 55 seconds with the following parameters: 450 peak power, 15% acoustic duty factor, 1000 cycles/burst, and 67.5 average power. The sheared DNA fragments were size selected by Solid PhaseReversible Immobilization using TotalPure Next Generation Sequencing (NGS) beads (Omega Bio-tek) and then the selected fragments were end-repaired, A-tailed, and ligated with Illumina compatible unique dual-index sequencing adaptors (IDT, Inc).

#### Sequencing

The prepared libraries were quantified using KAPA Biosystems’ next-generation sequencing library qPCR kit and run on a Roche LightCycler 480 real-time PCR instrument. Sequencing of the flowcell was performed on the Illumina NovaSeq sequencer using NovaSeq XP V1.5 reagent kits, S4 flowcell, following a 2×151 indexed run recipe. The Illumina NovaSeq 6000 instrument with the S4 type flow cell was used along with the Low Input DNA (400 bp fragment) library construction for sequencing. Reads were then quality controlled for length (>50 bp), quality (>Q30), and any adapter contamination was removed along with mateless pairs. Filtered reads were error-corrected using bbcms v.38.90^15^. This was run using the following command line options: bbcms.sh -Xmx100g metadatafile=counts.metadata.json mincount=2 highcountfraction=0.6 in=bbcms.input.fastq.gz out1=input.corr.left.fastq.gzout2=input.corr.right.fastq.gz. The readset was assembled with metaspades v. 3.15.2^16^. This was run using the following command line options: spades.py - m2000 --tmp-dir cromwell_root -o spades3 --only-assembler -k 33,55,77,99,127 --meta -t 16-1 input.corr.left.fastq.gz -2 input.corr.right.fastq.gz. The input read set was mapped to the final assembly and coverage information generated with bbmap v. BBMap:38.86^15^. This was run using the following command line options: bbmap.sh build=1 overwrite=true fastareadlen=500 - Xmx100g threads=16 nodisk=trueinterleaved=true ambiguous=random rgid=filename in=reads.fastq.gz ref=reference.fastaout=pairedMapped.bam. Following assembly, the metagenomes underwent processing through the DOE JGI Metagenome Annotation Pipeline (MAP version 5.1.13) and were subsequently loaded into the Integrated Microbial Genomes and Microbiomes (IMG/M) platform^17^. Complete metagenomic datasets for each time point are listed by individual IMG Genome IDs^12^ under the Study Name “Seawater microbial communities from East Sound, Orcas Island, WA, USA”. For each time point, estimated gene copies for all classes in the domain “Bacteria” were downloaded (3.14.2024; Dataset 5)^12^. At each time point, the cumulative count of gene copies was calculated, and subsequently, each taxonomic class was divided by the total count to determine the class distribution (Figure 4; Dataset 5)^12^.

**Figure 4.**
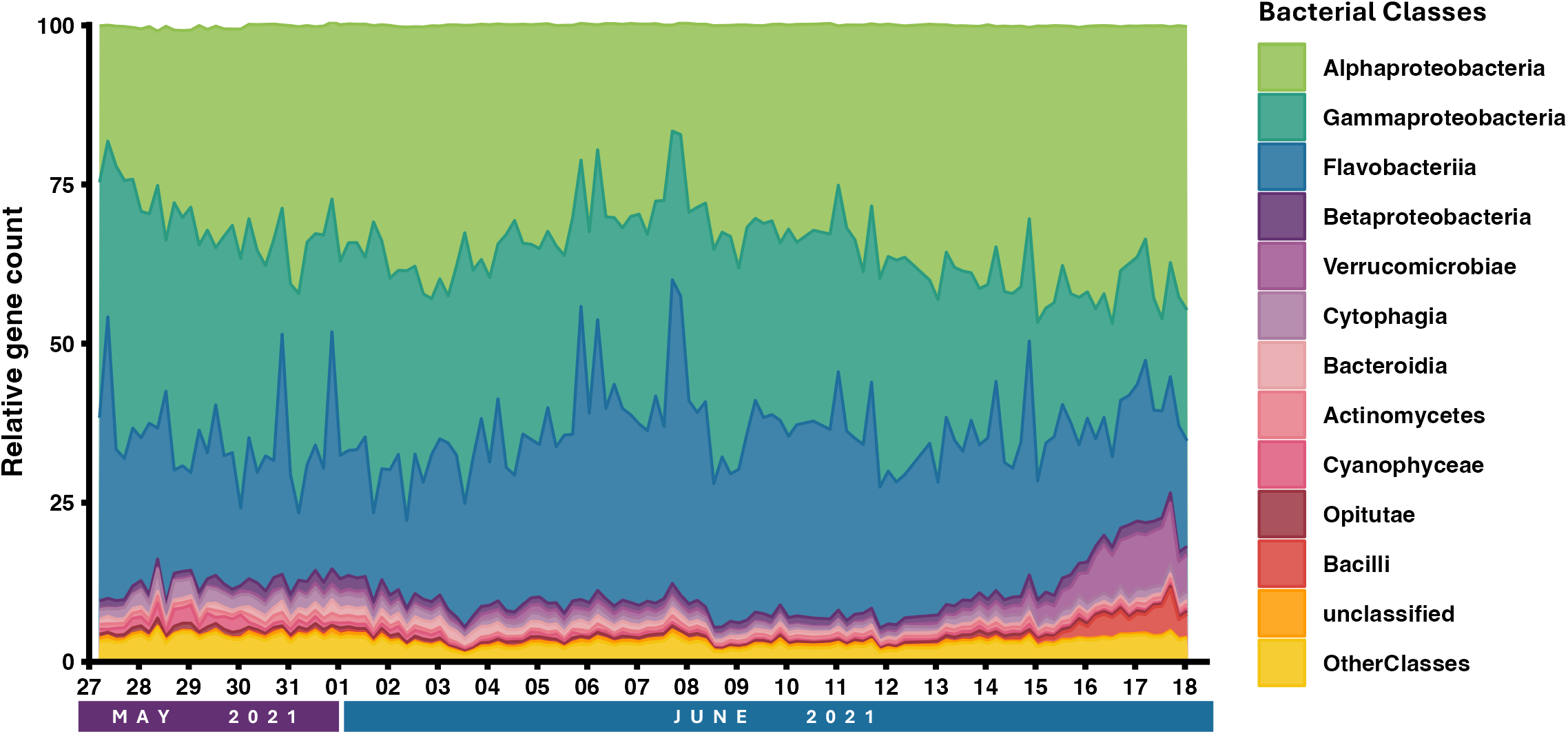
Relative abundance of bacterial classes in East Sound from May 27 13:00 - June 18 13:00 collected 6 times per day at 1:00, 5:00, 9:00, 13:00, 17:00, and 21:00. Distribution of bacterial classes based on estimated gene counts from metagenomes completed by JGI at each time point. Metagenomic samples were collected at 2 m depth in 2021 and microbiome community DNA from 0.22 to 1.0 □ *µ*m size range. Taxonomic groups were defined based on exact sequence variants and those without variants that were specified to a class were grouped together in the unclassified category (orange). Assignments of the 11 most abundant taxonomic classes are presented. Classes listed yielded averages > 0.5% of the total. All other classes not listed in the legend were summed as Other Classes.

### Data Records

The Illumina sequencing reads for metagenomes collected at each time point are individually accessible through the NCBI Sequence Read Archive, grouped under a single BioProject: PRJNA1093221 (https://www.ncbi.nlm.nih.gov/bioproject/PRJNA1093221)^12,18^. Metagenome-assembled genomes (MAGs) and contigs assembled at each of the 128 time points can be obtained from the JGI Integrated Microbial Genomes portal^12^ and under the Joint Genome Institute Genome OnLine Database^19^ (GOLD ID) Gs0160720. Datasets 1-5 can be accessed on figshare (https://doi.org/10.6084/m9.figshare.26882737)^12^. Details on the data contained in each file are as follows:

#### Dataset 1 - Orcas Island, WA, USA 2021 Coastal Ocean (2m depth) Time Series - Environmental YSI EXO1 Sonde Probe Data (Nunn_OrcasIsland_Data_Probe.xlsx)

Data collected by the YSI Probe includes columns for the date (Date) and time (Time) the sample was collected, a character value for the combined date and time of sample collection (Date.Time), chlorophyll relative fluorescence units (Chlorophyll (RFU)), chlorophyll *µ*gL (Chlorophyll (ug L-1)), electrical conductivity (Conductivity (us cm-1)), depth the sample was collected in meters (Depth.m), optical dissolved oxygen (ODO (% saturation)) and (ODO (mg L-1), salinity (Sal (psu)), pH (pH), the temperature in degrees Celsius (Temp (degrees Celsius),), and coordinates (Latitude and Longitude) of sample collection site.

#### Dataset 2 – Orcas Island, WA, USA 2021 Coastal Ocean (2m depth) Time Series - SoundToxins Phytoplankton Monitoring Network East Sound Report, Washington Sea Grant (Nunn_OrcasIsland_Data_SoundToxins.xlsx)

Data file consists of two tabs. Data includes relative abundance observations of phytoplankton at the SoundToxins East Sound Monitoring Site by citizen science volunteers. Note that all observational “Abundance” data is reported as categorical abundances based on microscopy and does not directly reflect the amount of biomass present in the water as represented by taxonomic group. Observations included for weekly site visits from 5/4/21 – 7/15/21. Credit: SoundToxins Phytoplankton Monitoring Network – Washington Sea Grant

- **EastSound_Visits:** Includes visit comments on environmental and water conditions and general observations of phytoplankton present in the water for the East Sound County Dock utilized by the SoundToxins program. Columns include site location (Name), Program, and Date/Time of sample collection. Columns detailing physical conditions at the site include Water, Air, Salinity, Depth Towed (m), Cod End, Wind, Weather, Tide, and Obs. General commentary for each visit date included in “Visit Comments” column.
- **EastSound_Observations:** Includes columns for Program (SoundToxins), Site (County Dock – East Sound), and date and time of observation (Date). For each Date, the phytoplankton observed in the sample are reported as Category, Genus and Species. Abundance measurements reflect the relative abundance of phytoplankton observed in sample as defined by SoundToxins Program (“Bloom” = dominant phytoplankton in sample, “Common” = phytoplankton abundant in sample, “Present” = phytoplankton observed in sample).

#### Dataset 3 – Orcas Island, WA, USA 2021 Coastal Ocean (2m depth) Time Series - Nutrient Analysis (Nunn_OrcasIsland_Data_Nutrients.xlsx)

Nutrient concentrations collected across time. Columns include the date the sample was collected (Date) and time of collection (Time), and a character value for the combined date and time of sample collection (DateTime). Phosphate concentrations (PO4 [mM]), silicate concentrations (Si.OH.4 [mM]), nitrate concentrations (NO3 [mM]), nitrate concentrations (NO2 [mM]), ammonium concentrations (NH4 [mM]), and coordinates (latitude and longitude) of sample collection site.

#### Dataset 4 – Orcas Island, WA, USA 2021 Coastal Ocean (2m depth) Time Series - Flow Cytometry Analysis (Nunn_OrcasIsland_Data_FlowCytometry.xlsx)

Columns for the day the sample was collected (1 of 22; Day), the date (Collection Date), and time (Hour) the sample was collected, a character value for the combined date and time of sample collection (DateID) are reported. Flow cytometry data includes cell counts (cells/mL) for triplicate analyses of cyanobacteria (Cyanobacteria Replicates 1-3), picoeukaryotes (Picoeukaryotes Replicates 1-3), nanoeukaryotes (Nanoeukaryotes Replicates 1-3), their averages and standard deviations. Bacteria were analyzed in duplicate and are included (Bacteria Replicates 1-2) with the average and standard deviation reported. Coordinates (latitude and longitude) of sample collection site are also included.

#### Dataset 5 – Orcas Island, WA, USA 2021 Coastal Ocean (2m depth) Time Series - Joint Genome Institute Metagenomic Sequencing (Nunn_OrcasIsland_Data_JGI_metadata): Data file consists of three tabs

- ***NCBI Sequence Read Archive IDs:*** Includes NCBI Sequence Read Archive ID numbers, grouped under a single BioProject: PRJNA1093221. Columns include the SRA Experiment Accession number, SRA Experiment Title (which indicates the location the sample was taken, the date the sample was taken, the day number the sample was taken out of 22 days, and the time the sample was taken 1:00, 5:00, 9:00, 13:00, 17:00, 21:00), instruments sequencing was completed on (Instrument), who submitted the archive entry (Submitter), study accession number, the study title, the sample accession number, the sample title, the total size of the file (Mb), the total number of runs, total spots, total bases, library name, library strategy, library source, library selection, and coordinates (latitude and longitude) of sample collection site.
- ***IMG Genome IDs:*** Complete list of JGI metagenomic datasets for each time point are listed by individual Integrated Microbial Genomes & Microbiomes (IMG) Genome IDs and include columns for the following parameters: JGI Integrated Microbial Genomes & Microbiomes identification number (IMG GenomeID), character value for the combined date and time of sample collection (DateID), type of sample sequenced (Domain), status of JGI sequencing effort for this ID (Sequencing Status), the name of the umbrella project study (Study Name), name of the specific sample includes standard annotation location_sample_datecollected_daycollected_timecollected (Genome Name / Sample Name), location of sequencing (Sequencing Center), Sample ID number (IMG Genome ID), Submission ID (IMG Submission ID), Joint Genome Institute Genome OnLine Database project ID (GOLD Analysis Project ID), type of Joint Genome Institute Genome OnLine Database project (GOLD Analysis Project Type), Joint Genome Institute Genome OnLine Database project ID (GOLD Sequencing Project ID), size of assembled genome (Genome Size * assembled), number og genese in metagenome (Gene Count * assembled), number of metaBAT counts, (Genome MetaBAT Bin Count * assembled), estimated number of genomes assembled (Estimated Number of Genomes * assembled), estmated average genome size (Estimated Average Genome Size * assembled).
- ***Estimated Gene Copy Number:*** The Estimated gene copies for all classes in the domain “Bacteria” for each time point. Data downloaded from JGI 3.14.2024. The first column is the JGI sample ID for the specified timepoint (IMG Genome ID) followed by a character value for the combined date and time of sample collection (DateID) and all the classes identified by JGI with their respective estimated gene copy numbers.

All R code (version 4.4.0) used for analyses are available on the GitHub page associated with this project: https://github.com/Nunn-Lab/Publication-2021-Orcas-Island-Time-Series

### Technical Validation

For Illumina metagenomes, BBDuk (version 39.03) was used to remove contaminants, trim reads that contained adapter sequence and homopolymers of G’s of size 5 or more at the ends of the reads, and right quality trim reads where quality drops to 0. BBDuk was also used to remove reads that contained 4 or more ‘N’ bases, had an average quality score across the read less than 3, or had a minimum length <= 51 bp or 33% of the full read length. Reads mapped with BBMap to masked human, cat, dog, and mouse references at 93% identity were separated into a chaff file. Reads aligned to common microbial contaminants were separated into a chaff file. Additional details on data handling, assembly, and annotation can be found in Clum et al., 2021^17^.

### Usage Notes

Metagenome data can be accessed via IMG/M using the Advanced query search: JGI GOLD IMG IDs -- ITS Proposal ID 509101. Data can be accessed via JGI’s Genome Portal (https://genome.jgi.doe.gov/portal/Invtheabustcycle/Invtheabustcycle.info.html).

## Code Availability

The workflow modules for read filtering, quality control, and metagenome assembly along with Cromwell workflow manager information are available as a workflow description language (WDL) file at https://code.jgi.doe.gov/BFoster/jgi_meta_wdl. The IMG/M annotation workflow is available at https://code.jgi.doe.gov/official-jgi-workflows/jgi-wdl-pipelines/img-omics. All R code used for analyses is available on the GitHub page associated with this project: https://github.com/Nunn-Lab/Publication-2021-Orcas-Island-Time-Series

## Supporting information

Nunn_OrcasIsland_Data_FlowCytometry.xlsx

Nunn_OrcasIsland_Data_JGI_metadata

Dataset 2: Nunn_OrcasIsland_Data_Nutrients.xlsx

Nunn_OrcasIsland_Data_Probe.xlsx

Nunn_OrcasIsland_Data_SoundToxins.xlsx

DataFile_Descriptions

## Acknowledgment

We would like to thank the 2021 field team for their Herculean efforts of collecting samples 24 hours a day for 22 days. In addition to the author list, this included Maya Garber-Yonts, Kathryn Sharp, Edith Branner, Katherine Riffle, Sam Hart, Kusum Dhakar, and Laura Gonzalez. The fieldwork would not have been possible without permission provided by the Youngren Family to use their dock and home for fieldwork, and Long Live the King’s Hatchery and Michael O’Connell for the use of tools, property, and general friendliness. UW Oceanography Technology provided the use of ropes and buoys to build the temporary mooring site and Angela and Stuart Currie provided tugboat support throughout the field expedition. Ardi Kveven and Katherine Dye provided the use of their YSI probe for the duration of the project in addition to calibration standards. Fieldwork was funded by a pilot grant from The University of Washington Royal Research Foundation to BL Nunn in 2021 and donations from the Nunn family. Support for personnel was provided by NIH grant R21ES034337-01 to BL Nunn and J Kubanek and NSF IOS grant NSF IOS 2041497 to BL Nunn. The work (proposal: DOI 10.46936/10.25585/60008633) conducted by the U.S. Department of Energy Joint Genome Institute (https://ror.org/04xm1d337), a DOE Office of Science User Facility, is supported by the Office of Science of the U.S. Department of Energy operated under Contract No. DE-AC02-05CH11231.

## Author Contributions

BL Nunn, E Timmins-Schiffman, MC Mudge, G Chebli, J Kubanek, and W Noble designed and carried out the fieldwork, sample collections, and data analysis. MC Mudge and E Timmins-Schiffman performed all DNA extractions. M Riffle, D Plubell, and T Nunn performed fieldwork and sample collection. T Nunn provided metagenomic class-level distribution analysis and figure generation. E Harvey performed flow cytometry analyses. The entire JGI team performed metagenomic sequencing, data analysis, and web-based data distribution. T Glavina del Rio was the JGI project manager and liaison between JGI and BL Nunn. E Eloe-Fadrosh contributed to writing the methods. BL Nunn was the primary author of the manuscript and all others contributed to editing.

## References

1 Alkire, M. B. et al. Estimates of net community production and export using high-resolution, Lagrangian measurements of O2, NO3−, and POC through the evolution of a spring diatom bloom in the North Atlantic. Deep Sea Research Part I: Oceanographic Research Papers 64, 157–174 (2012).

2 Chitkara, C. et al. Seasonality in phytoplankton communities and production in three Arctic fjords across a climate gradient. Progress in Oceanography 227, 103317 (2024).

3 Henson, S., Le Moigne, F. & Giering, S. Drivers of carbon export efficiency in the global ocean. Global biogeochemical cycles 33, 891–903 (2019).

4 Sverdrup, H. On conditions for the vernal blooming of phytoplankton. J. Cons. Int. Explor. Mer 18, 287–295 (1953).

5 Buchan, A., LeCleir, G. R., Gulvik, C. A. & González, J. M. Master recyclers: features and functions of bacteria associated with phytoplankton blooms. Nature reviews. Microbiology 12, 686–698 (2014).

6 Cirri, E. & Pohnert, G. Algae− bacteria interactions that balance the planktonic microbiome. New Phytol 223, 100–106 (2019).

7 Needham, D. M. & Fuhrman, J. A. Pronounced daily succession of phytoplankton, archaea and bacteria following a spring bloom. Nat Microbiol 1, 16005, doi:10.1038/nmicrobiol.2016.5 (2016).

8 Li, W. & Dickie, P. Monitoring phytoplankton, bacterioplankton, and virioplankton in a coastal inlet (Bedford Basin) by flow cytometry. Cytometry: The Journal of the International Society for Analytical Cytology 44, 236–246 (2001).

9 Nowinski, B. et al. Microbial metagenomes and metatranscriptomes during a coastal phytoplankton bloom. Scientific data 6, 129 (2019).

10 Muratore, D. et al. Complex marine microbial communities partition metabolism of scarce resources over the diel cycle. Nature Ecology & Evolution 6, 218–229 (2022).

11 NOAA. NOAA Tide Predictions: 9449771 Rosario, Orcas Island, WA <https://tidesandcurrents.noaa.gov/noaatidepredictions.html?id=9449771&units=standard&bdate=20210501&edate=20210531&timezone=LST/LDT&clock=12hour&datum=MLLW&interval=hilo&action=dailychart> (

12 Nunn, B. L. et al. Microbial Metagenomes Across a Complete Phytoplankton Bloom Cycle: High-Resolution Sampling Every 4 Hours Over 22 Days, 2024).

13 UNESCO. Protocols for the Joint Global Ocean Flux Study (JGOFS) Core Measurements. 29 (1994).

14 Valderrama, J. C. The simultaneous analysis of total nitrogen and total phosphorus in natural waters. Mar Chem 10, 109–122 (1981).

15 Bushnell, B. BBTools software packag. e (2014).

16 Nurk, S., Meleshko, D., Korobeynikov, A. & Pevzner, P. A. metaSPAdes: a new versatile metagenomic assembler. Genome Res 27, 824–834 (2017).

17 Clum, A. et al. DOE JGI metagenome workflow. Msystems 6, 10.1128/msystems.00804-00820 (2021).

18 EMBL-EBI ENA browser, <https://identifiers.org/bioproject:PRJNA1093221> (2024).

19 Mukherjee, S. et al. Twenty-five years of Genomes OnLine Database (GOLD): data updates and new features in v. 9. Nucleic acids research 51, D957–D963 (2023).

20 Lawrenz, E. & Richardson, T. L. How does the species used for calibration affect chlorophyll a measurements by in situ fluorometry? Estuaries and Coasts 34, 872–883 (2011).

