## Supplementary material for "Microbial Metagenomes Across a Complete Phytoplankton Bloom Cycle: High-Resolution Sampling Every 4 Hours Over 22 Days": DataFile_Descriptions

**Nunn_OrcasIsland_Data_Probe.xlsx:** Data collected by the YSI Probe includes columns for the date (Date) and time (Time) the sample was collected, a character value for the combined date and time of sample collection (Date.Time), chlorophyll relative fluorescence units (Chlorophyll.RFU), chlorophyll µgL (Chlorophyll.ugL), electrical conductivity (Conductivity.uscm), depth the sample was collected in meters (Depth.m), optical dissolved oxygen (ODO.sat) and (ODO.mgL), salinity (Sal.psu), pH (pH), the temperature in degrees Celsius (Temp.C),), and coordinates (latitude and longitude) of sample collection site.

**Dataset 2: Nunn_OrcasIsland_Data_Nutrients.xlsx** Nutrient concentrations collected across time. Columns include the date the sample was collected (Date) and time of collection(Time), and a character value for the combined date and time of sample collection (DateTime). Phosphate concentrations (PO4 [mM]), silicate concentrations (Si.OH.4 [mM]), nitrate concentrations (NO3 [mM]), nitrate concentrations (NO2 [mM]), ammonium concentrations (NH4 [mM]), and coordinates (latitude and longitude) of sample collection site.

**Nunn_OrcasIsland_Data_FlowCytometry.xlsx:** Columns for the day the sample was collected (1 of 22; Day), the date (Collection Date), and time (Hour) the sample was collected, a character value for the combined date and time of sample collection (DateID) are reported. Flow cytometry data includes cell counts (cells/mL) for triplicate analyses of cyanobacteria (Cyanobacteria Replicates 1-3), picoeukaryotes (Picoeukaryotes Replicates 1-3), nanoeukaryotes (Nanoeukaryotes Replicates 1-3), their averages and standard deviations. Bacteria were analyzed in duplicate and are included (Bacteria Replicates 1-2) with the average and standard deviation reported. Coordinates (latitude and longitude) of sample collection site are also included.

**Nunn_OrcasIsland_Data_SoundToxins.xlsx:** Data includes relative abundance measures of phytoplankton at the SoundToxins East Sound Monitoring Site by citizen science volunteers. Credit: SoundToxins Phytoplankton Monitoring Network – Washington Sea Grant. Data file consists of two tabs. Note that all data is reported as relative abundance measures based on microscopy and does not reflect the amount of biomass present in the water. Observations included for weekly site visits from 5/4/21 – 7/15/21.
