## Supplemental Figure 1 for "Microbial Metagenomes Across a Complete Phytoplankton Bloom Cycle: High-Resolution Sampling Every 4 Hours Over 22 Days"

**Table of Contents	Page Number**

Supplemental Figure 1……………………………………………………………………………….3-4

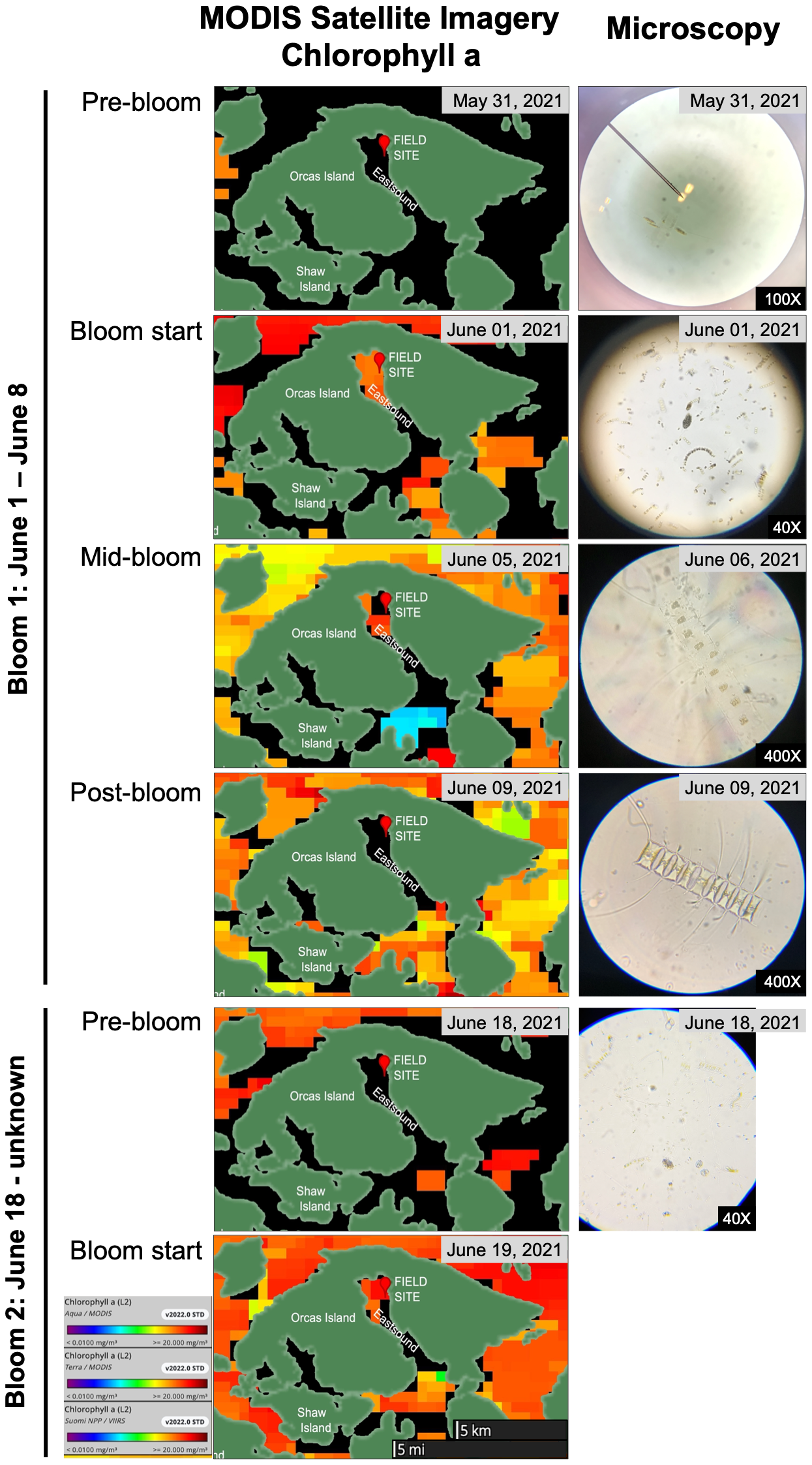

**Supplemental Figure 1. MODIS satellite imagery and on-site microscopy confirm the presence of two distinct phytoplankton blooms.** Representative satellite and microscopic images confirm pre-bloom and bloom start for two phytoplankton blooms, as well as mid-bloom and post-bloom of the first phytoplankton bloom. MODIS satellite images of near-surface chlorophyll a concentration (mg m-3) in waters surrounding East Sound, WA, USA. The MODIS Chlorophyll a product is available from both the Terra and Aqua satellites. The sensor and imagery resolution is 1 km, and the temporal resolution is daily. References: MODISA_L2_OC doi:10.5067/AQUA/MODIS/L2/OC/2018. Microscopy images of whole water collected on-site confirm presence of *Chaetoceros spp.* throughout field season.
